# Comparative study of Dog and Human Lipid Droplet-associated protein (PLIN2)

**DOI:** 10.1101/2025.11.17.688867

**Authors:** Ghanshyam Sahu, Amrita Behera, Vineet Kumar Pandey, P.S. Franco, Mukesh Kumar, Mohini Saini, I. Karuna Devi

## Abstract

Intracellular lipid droplets (LDs) are vital organelles that store energy, manage oxidative stress, and coordinate cellular signaling. A key protein family associated with LDs is the perilipins, with perilipin 2 (PLIN2) being prominent. This study focuses on a comparative analysis of canine and human PLIN2 to understand their structural and functional similarities. The genomic data for PLIN2 in both species were curated, revealing location, transcripts, exons, and introns. Sequence analysis demonstrated 85% identity between dog and human PLIN2, with conserved residues essential for structural and functional roles. Amino acid composition analysis revealed similarities in charged residues and in stabilizing and destabilizing residues. Despite their structural similarities (72% common secondary structure elements), a 3D comparison revealed significant differences in conformation. Physicochemical properties, including stability indices, were found to be distinct, possibly influencing their interactions. The study highlights the conservation of PLIN2’s functional regions, shedding light on its role in protecting LDs from degradation across species.

## Introduction

Intracellular lipid droplets (LDs) are lipid organelles that serve as a reservoir for energy sources, a sink for reactive oxygen species, and a hub for several cellular signals. Excessive degradation of these LDs can alter the cellular metabolism as well as pose threats for lipotoxicity and many other metabolic disorders (Olzmann and Carvalho, 2019). LDs consist of core forms by cholesterol ester, triacylglycerol that are surrounded by phospholipid monolayers, containing various proteins, and perilipin proteins are the most dominant structural proteins associated with LDs. Perilipin proteins are involved in the storage and protection of intracellular lipid droplets from cytosolic lipolytic lipases as well as many cellular signaling processes (Kuniyoshi et al., 2019).

Perilipin family consisting of five member namely perilipin1 (PLIN1), perilipin2 (PLIN2), perilipin3 (PLIN3),perilipin4(PLIN4) and perilipin5 (PLIN5). PLIN2 is the major perilipin which is found ubiquitous and expressed widely in hepatocytes, mammary gland, adipose tissues, foam cells, sebocytes (Itabe et al., 2017). PLIN2 plays an important role in governing LDs dynamics and therefore, PLIN2 is chiefly focused in this paper for detailed investigations.In the current manuscript, we have performed the comparative study of Dog and Human PLIN2 for their structural and functional similarities using the sequences and structure related data (Griseti et al., 2024). In our study, we found 85% sequence similarity and 72% structural similarity in between Dog and Human PLIN2. However, in the majority of the cases, we could not see significant differences in the structural properties and amino acid compositions between them which suggest that these proteins have similar roles in protecting the LDs degradation.

## Materials and Methods

1. **Data curations:** We performed manual curation of data associated with PLIN2 genes, transcripts and proteins. The Perillipine 2 genes and protein-related information of two species (Human and Dog) were retrieved from the NCBI (Sayers et al., 2025) and Uniprot (“UniProt: the universal protein knowledgebase in 2025,” 2025) respectively. Similarly the 3D structure of dog (AF-E2RP20-F1-model_v4) and human PLIN2 (AF-Q99541-F1-model_v4) proteins were retrieved from AlphaFold database(Fleming et al., 2025).
2. **Multiple sequence alignments (MSA):** We performed MSA of perilipin proteins belongs to five different species using the CLUSTAL OMEGA web server.
3. **Protein compositions and physicochemical properties analysis:** The Expasy tool “ProtParam” was employed to analyze the protein compositions and physicochemical properties of Dog and Human PLIN2 proteins i.e, amino acid composition, molecular weight, instability index, theoretical isoelectric point (pI), aliphatic index, and grand average of hydropathicity (GRAVY).
4. **Secondary structure prediction and analysis:** The secondary structure of Dog and Human PLIN2 proteins is predicted and analyzed using the DSSP and Bio3D R package (Al-Azzawi and Alsaedi, 2025).
5. **Secondary structure analysis:** DALI web server was used to compare the secondary structures between Dog and Human PLIN2 proteins (Holm, 2022).
6. **Protein structure quality analysis:** it is done by using VADAR (Volume, Area, Dihedral Angle Reporter) server(Anand et al., 2025).

## Results and Discussion

### a. Genomics Transcript Variants of PLIN2

The most recent canine (**Dog10K_Boxer_Tasha**) and human (**GRCh38.p14**) reference genome assemblies were used in the curation of the Dog **PLIN2** (Gene ID: 100856329) and Human **PLIN2** (Gene ID: 123) genomic information obtained from the NCBI and UCSC Genome Browser databases (Goldfarb et al., 2025; Jagannathan et al., 2021) Dog PLIN2 is located on chromosome 11, in the region between 37598800 and 37617859 nucleotides (Location ID: NC_006593.4). Similar to its canine counterpart, the Human PLIN2 gene is situated on chromosome 9 specifically in the short arm, second region, second band, and first sub band between 19108388 and 19127492 nucleotides (Location ID: NC_000009.12, (9p22.1). Dog PLIN2 has eleven exons, according to the NCBI gene annotation, although we could not see the same numbers in the UCSC genome browser. So, taking into account 10 exons that can segregate differentially to create partial and full forms of transcripts (XM_003639380.5, XM_005626663.), and subsequently matching proteins (Fig.1). Similarly, human PLIN2, which has a maximum of eight exons generates three transcript variants (NM_001122.4, NR_038064.2, and XM_017014259.3) each of which can result in the production of either partial or complete proteins. NCBI gene annotation shows eleven exons in Dog PLIN2 but we could not validate it in UCSC genome annotations. So, considering 10 exon that can segregate differently to form multiple transcript variants and produce partial and full forms of transcripts (**XM_038681839.1, XM_038681837.1, XM_038681838.1**) and subsequently proteins (Fig.1). Likewise, the maximum number of exons in human PLIN2 is eight that gives three transcript variants (**NM_001122.4**,**NR_038064.2**,**XM_017014259.3)** and produce partial and full proteins. Different proportions of exons are incorporated during the transcripts and protein synthesis in different pathophysiological conditions. These variants could be unregulated or down-regulated in context dependent manner as well as such exonic variation may affect the structural and functional properties of the PLIN2 proteins.

**Fig 1:**
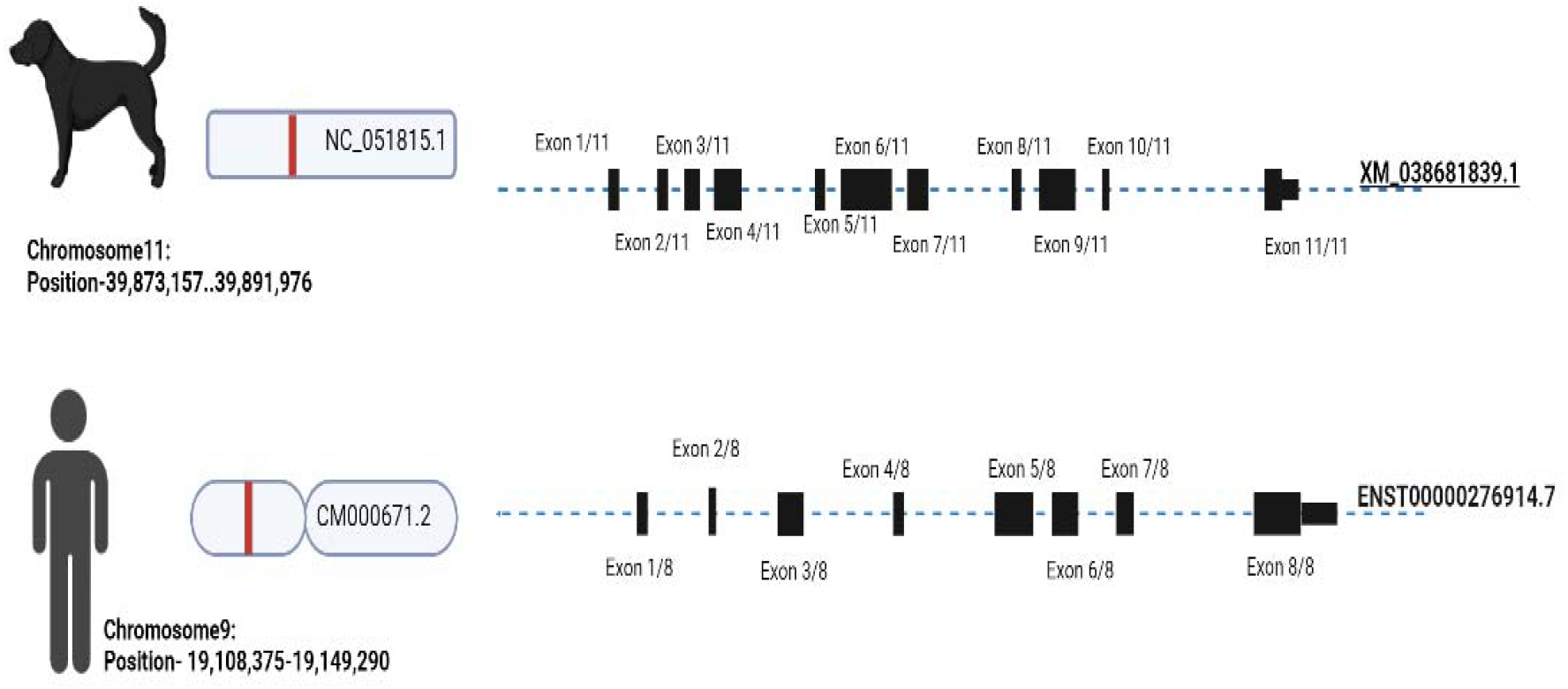
Chromosomal location and transcript variants of canine PLIN2 gene.

**Fig 2.**
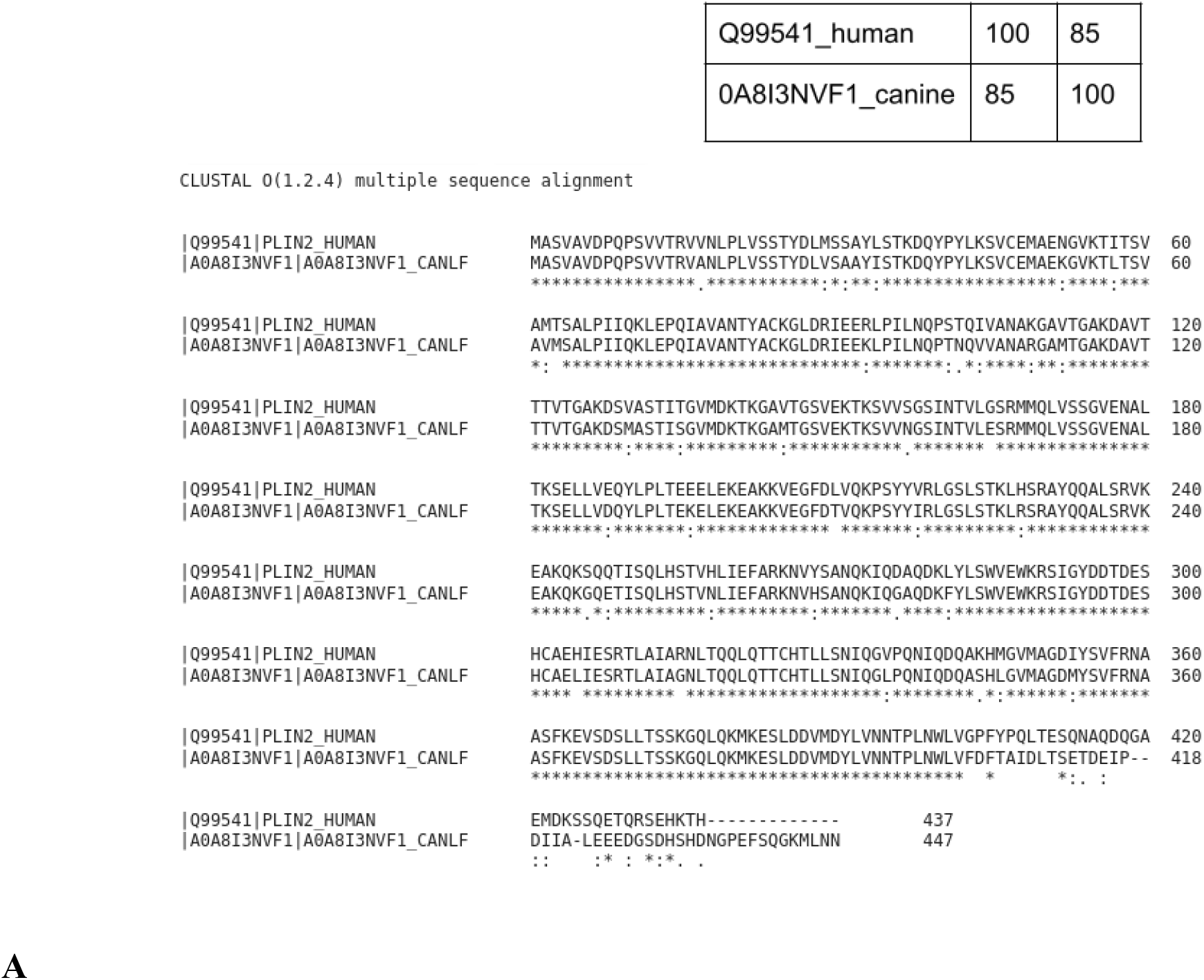
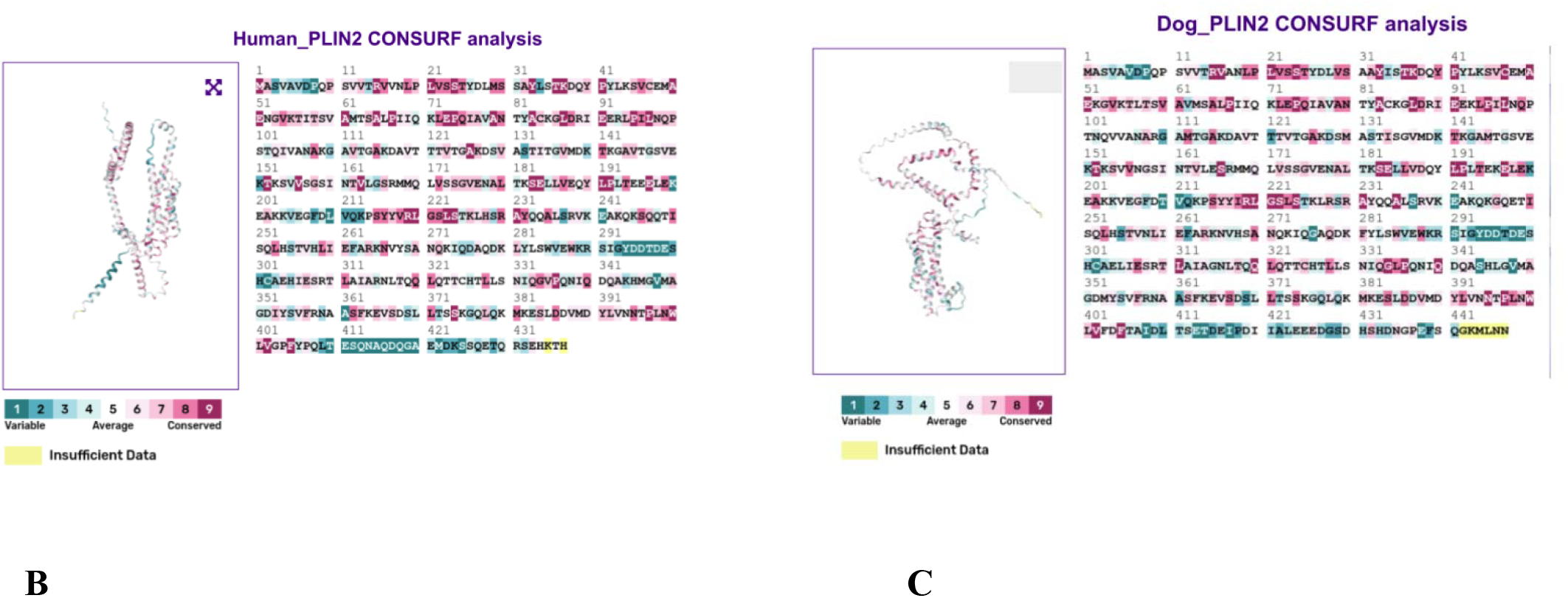
Sequence similarity and evolutionary conservation of canine and human PLIN2 proteins.(Fig.2A) Multiple sequence alignment of canine PLIN2 (A0A8I3NVF1) and human PLIN2 (Q99541) generated using CLUSTAL Omega, showing high sequence identity (85%) and conserved functional regions.**(Fig2B–2C)** ConSurf analysis illustrating residue-level evolutionary conservation mapped onto the predicted 3D structure and linear sequence of Human (B) and Dog(C)PLIN2. Highly conserve residues (purple) correspond to functionally or structurally important regions, whereas variable residues (cyan) indicate species-specific divergence; yellow denotes positions with insufficient data.

**Fig 3.**
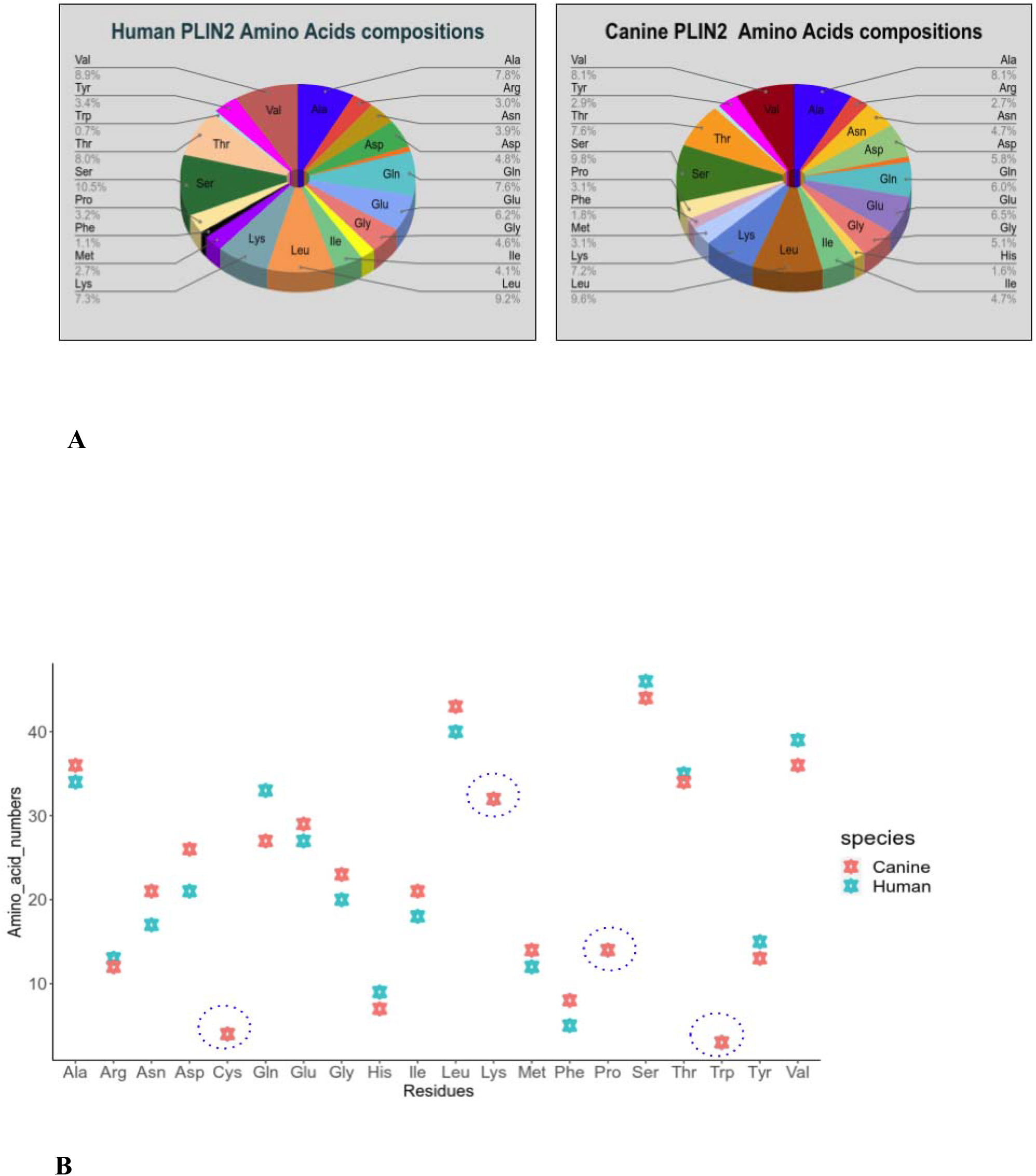
Amino acid compositions and physicochemical properties of PLIN2 proteins. Fig.3A & Fig.3B shows the differences of amino acid compositions (percentages and numbers) in Human and Dog PLIN2 proteins. Encircled residues are equally present in both species.

**Fig 4:**
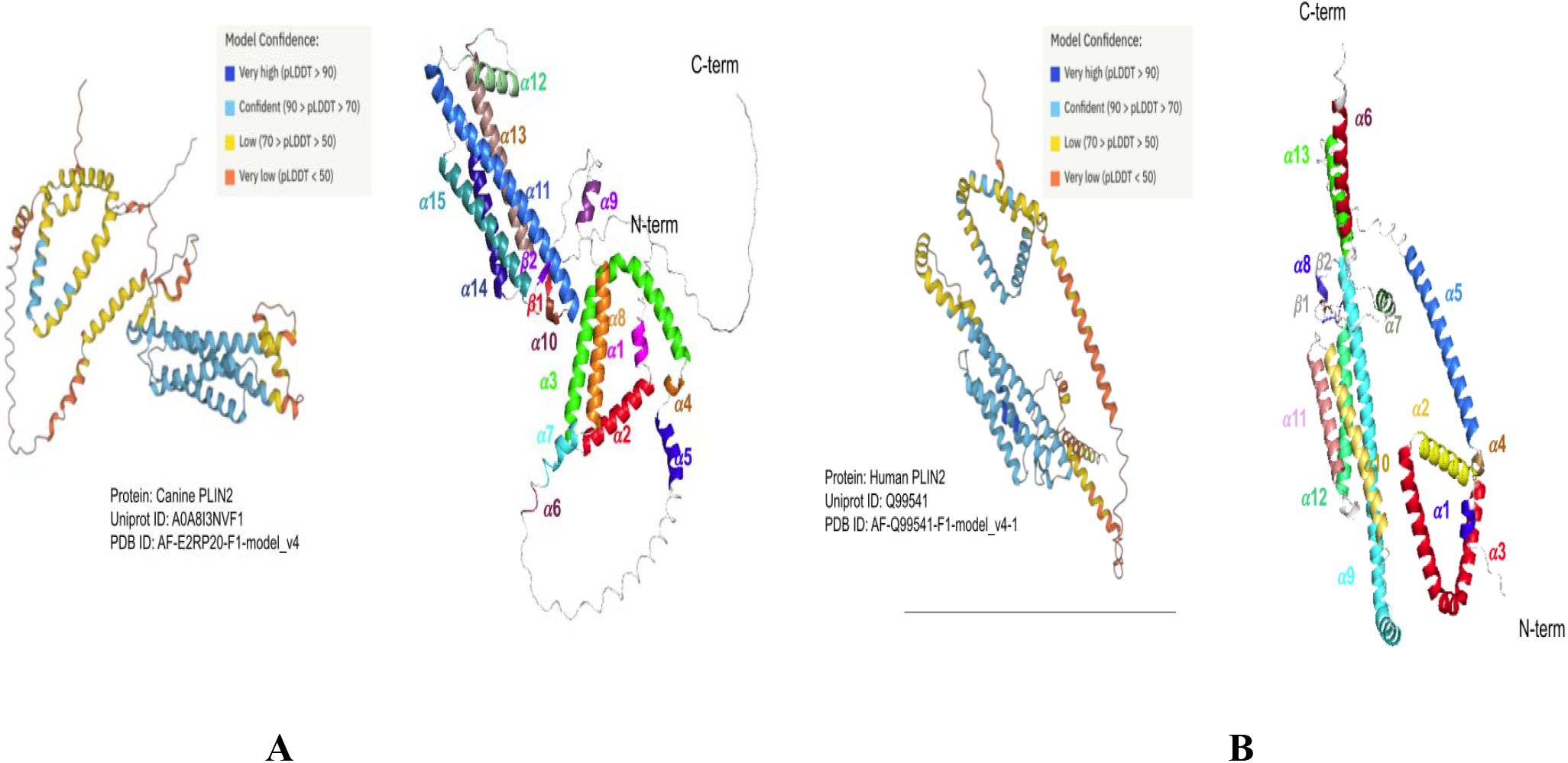
3D structure elements of PLIN2, which is based on the analysis and annotation using Bio3D R package,(**Fig.4A)** showed the secondary structure elements of Human,**(Fig.4B)** showed the secondary structure elements of canine

### b. Sequence similarity and residues conservation analysis

The amino acid sequences of Human and Dog Perilipin 2 (PLIN2) proteins are retrieved from the Uniprot database (“UniProt: the universal protein knowledgebase in 2025,” 2025) and compared for percent identity and conserved residues using the CLUSTAL OMEGA webserver. Such study could be relevant for structural and functional similarity of this protein among different species. We found Dog PLIN2 (447 amino acid) is a little longer and 85 percent identical to the human counterpart (437 amino acid). The C-terminal of Dog PLIN2 is less conserved. Using the ConSurf server, we further explored the Dog and Human PLIN2 for sequence conservation and its significance for functional regions in the proteins using the CONSURF server. CONSURF analysis replicates the results of the CLUSTAL omega where one can see that the majority of the residues (except C-terminus) are quite conserved in both proteins. Such residue conservation among the species shows that these proteins probably possess structural and functional similarity to protect the lipid droplets from lipolysis cytosolic lipases in two different species.

## Percent identity matrix

### c. Physicochemical properties of PLIN2 proteins

Diversities in amino acid composition play an important role for maintaining the stability and structural and functional specificities. Comparing the amino acid compositions between dog and human PLIN2 suggests how much these proteins behave differently on lipid droplet interactions. The positively (45,44) and negatively (48,55) charged residues are significantly abundant in both canine and human PLIN2 proteins. Presence of charge residues could be the player for major non-covalent interactions such as hydrogen bond, electrostatic interaction and salt bridge. Such kinds of interactions are pivotal not only for interactions with lipid droplets but also for many allosteric interaction where long-range communications are involved (chiefly by electrostatic interactions). Both PLIN2 contain identical quantities of the four residues Cys, Lys, Pro, and Trp despite the larger length of Dog PLIN2 protein. Cysteine can be involved in disulfide bonding; likewise electrostatic interactions can be facilitated by Lysine. The presence of Tryptophan can enlarge the hydrophobic surface due to bulkiness and larger surface area. Human PLIN2 is more stable than canine PLIN2 based on the instability index, which is 42.63 for canine PLIN2 and 37.24 for human PLIN2. The estimated half-life of these proteins, which is 30 hours for both proteins, is unaffected by these differences.Such differences could be affected due to variation in th numbers of polar residues, instability or stability inducing residues (PMID: **2075190**)

We found the number of polar (P)residues (P: “Arg”,”Asn”, “Asp”, “Cys”,”Gln”, “Glu”, “His”, “Lys”,”ser”, “Thr”, “Tyr”) and instability inducing residues (IIR) (“Met”,”Gln”, “Glu”, “ser”, “Pro) are quite same in dog and human PLIN2 proteins that is (P; Dog:205, Human 206 aa) and (IIR; Dog:84 and Human 86 aa) respectively. However surprisingly, the stability inducing residues (SIR) are quite low in Human PLIN2 (76 in Dog vs 69 in Human PLIN2), despite it, human PLIN2 shows higher instability due to unknown reasons that needs to be resolved in future.

### d. Data curation and annotation of three dimensional (3D) structure of PLIN2

Experimental 3D structures of PLIN2 are not available in protein data banks (PDB)

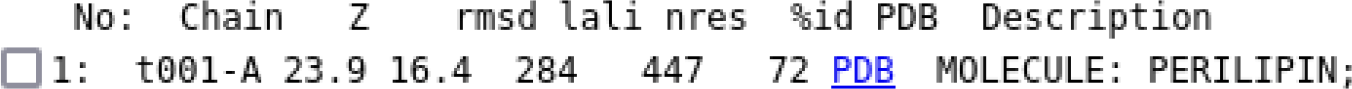

(Ref: PMID: **36419248**). However, homology modeling has been employed in numerous studies to forecast the 3D structure, but it has been known for being inaccurate. The AlphaFold server recently predicted the 3D structures of many proteins using neural network-based machine learning algorithms. These predictions were determined to be particularly promising for some of the proteins and parallel to the experimental structures after passing the Critical Assessment of Protein Structure Prediction14 (CASP14) test (Fleming et al., 2025). By Considering these advantages, we retrieved the human (AF-Q99541-F1-model_v4) and canine PLIN2 3D structures (AF-E2RP20-F1-model_v4_canine). The confidence level of structure prediction was shown as such available in the original source i.e. alphaFold database. These structures are of moderate quality in which some of the regions show low confidence scores and could be unstructured. The discrepancies between the canine and human PLIN2 conformations can be examined using RMSD. We compared the canine and human PLIN2 using the DALI web server (Holm, 2022), and discovered relatively large values of RMSD (>16). These results showed that both PLIN2 are structurally diverse despite being very similar in the secondary structure contents (72%) and amino acid compositions. The details of the structure alignment is shown below.

### e. Secondary structure analysis

We used the standalone version of the DSSP program as well as the Bio3D R package to analyze the secondary structure contents in both Human and Canine PLIN2 (Grant et al., 2006). We found canine PLIN2 consists of 15 *α*-helices, 2 *β*-sheets and 12 *β*-turns. The *α*-helices are *α*-helix-1 (12-17), *α*-helix-2(20-39), *α*-helix-3 (41-93), *α*-helix-4 (95-98), *α*-helix-5 (101-112), *α*-helix-6 (150-152), *α*-helix-7 (155-164), *α*-helix-8(167-190), *α*-helix-9 (195-201), *α*-helix-10 (220-222), *α*-helix-11 (225-273), *α*-helix-12 (279-291), *α*-helix-13 (299-331), *α*-helix-14 (337-359), *α*-helix-15 (368-394). The two *β*1 and *β*2 sheets expand in between 217-219 and 401-403, respectively. In alpha-helices dominant PLIN2, Helix-3 is the longest, followed by helices 14 and 15, which are the shortest. Likewise, Human PLIN2 consists of 12 *α*-helices, 2 *β*-sheets and 10 *β*-turns. The *α*-helices of human PLIN2 are *α*-helix-1 (12-17), *α*-helix-2 (20-39), *α*-helix3 (41-93), *α*-helix-4 (95-98), *α*-helix-5 (101-137), *α*-helix-6 (167-188), *α*-helix-7 (195-201), *α*-helix-8 (220-222). *α*-helix-9 (225-292), *α*-helix-10 (300-331), *α*-helix-11 (337-359), *α*-helix-12 (368-394), *α*-helix-13 (413-431). The two *β*1 and *β*2 sheets expand in between 216-219 and 401-405, respectively.In alpha-helices dominant PLIN2, Helix-7 is the longest, followed by helices 12 and 13, which are the shortest (Al-Azzawi and Alsaedi, 2025).

### f. Comparison between Human and Canine PLIN2 3D structure quality

We used VADAR (Volume, Area, Dihedral Angle Reporter) to predict the quantitative and qualitative assessment of structural properties of canine PLIN2 and compared them with the human PLIN2 structure (Willard et al., 2003). We examine the Ramachandran plot for secondary structures (Fig.5A), Backbone angle (Fig.5B), solvent accessibility (Fig.5C), Residual fractional Volume (Fig.5D), and packing quality and intrachain hydrogen bonds. Human PLIN2 has a better distribution of secondary structures than canine PLIN2. Canine PLIN2 has a lot more secondary structures in the first quadrant of the Ramachandran plot, which is the known location for left-handed helices and where most of them are in forbidden areas. Solvent accessible surface analysis in canine and human PLIN2 shows no significant differences. In both species, we found that the majority of the residues (> 370 residues) have high solvent accessibility, considering the >30 as a cutoff for accessibility. Less than 30 was considered buried. Similarly, we have observed quite similar residue functional volume in both PLIN2 structures. However, the extreme c-terminal end of canine PLIN2 shows a poor score (less than 0.8) of fraction volume which indicates that this region can suffer from packing defects. The distributions of ⍰ (Omega) angles are also analyzed and found that significant proportions of c-terminal residues of canine PLIN2 fall below 170^0^ compared to human PLIN2 This indicates that these residues could have distorted backbone angles in Canine PLIN2.

**Figure 5.**
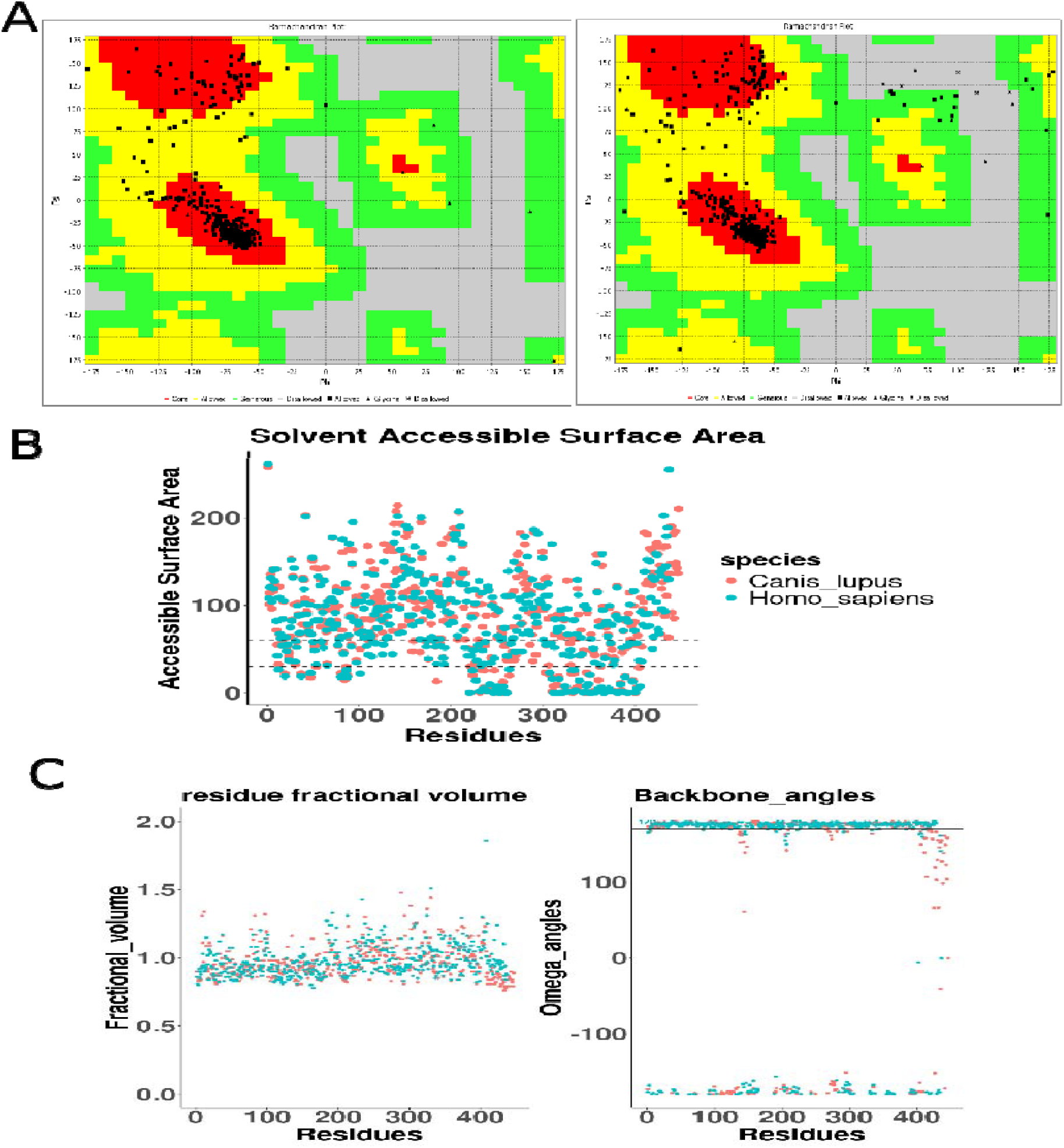
Protein structure quality assessment and comparative residue-level properties of Human and Canine PLIN2.(Fig.5A) Ramachandran plot analysis of Human (left) and Canine (right) PLIN2 structures. The plots display the distribution of backbone φ/ψ dihedral angles. Most residues fall within favored and allowed regions, indicating high stereochemical quality for both models. Minor deviations in disallowed regions reflect species-specific loop flexibility.**(Fig.5B)** Solvent Accessible Surface Area (SASA) comparison between Human (cyan) and Canine (salmon) PLIN2. Scatter plot shows residue-wise SASA distribution across the full sequence. Dashed lines mark thresholds commonly associated with buried (<30 Å^2^) and partially exposed (∼60 Å^2^) residues, illustrating broadly conserved surface exposure patterns between species with localized variations.**(Fig.5C)** Residue fractional volume and backbone ω-angle comparison.**Left panel:** Fractional volume distribution of Human and Canine PLIN2 residues, showing overall conservation of residue packing density with isolated variations in bulky or flexible regions. **Right panel:** Backbone ω-angle distribution highlighting trans-peptide bond predominance (∼180°) for both species, with minor deviations corresponding to flexible or strained regions.

## Solvent accessibility analysis (SASA)

Both structures of canine and human PLIN2 were subjects in the VADAR webserver in order to analyze the SASA, where the figure shows 0.30 was considered as buried, >0.30 to <0.60 wa considered as moderately solvent exposed and >0.60 was considered as highly solvent exposed residues.

## Conclusion and future perspective

### Conclusion

In this study, a comprehensive analysis of canine and human perilipin 2 proteins was conducted to investigate their structural and functional similarities. Despite an 85% sequence identity and 72% common secondary structure elements, significant structural differences were observed in their 3D conformations. Important functional areas were preserved across species, as evidenced by the identification of conserved residues essential for functional tasks. Protein-lipid interactions may be impacted by differences in charge residues, stabilizing residues, and destabilizing residues, which were identified by physicochemical property study. The results provide a basis for additional research into the molecular processes controlling cellular signaling, lipid metabolism, and possible treatments for metabolic diseases.

### Future Directions

This work offers a fundamental framework for understanding perilipin 2’s intricate biological functions in lipid droplet biology and general cellular homeostasis. Even though the present results provide valuable comparative insights, there are still several untapped study directions that need thorough exploration.

#### Functional Validation

To confirm the anticipated functional roles of conserved residues and how they support cellular signaling and LD protection, experimental research is required.

#### Structural Dynamics

Gaining a greater understanding of perilipin 2’s functionality can be achieved by comprehending its dynamic activity in response to different cellular stimuli and interactions with lipids.

#### Species Variability

By extending this investigation to additional species, it may be possible to identify evolutionary patterns in the conservation and divergence of perilipin 2, providing insight into species-specific adaptations.

#### Pathophysiological Significance

Researching how structural and functional differences in perilipin 2 affect lipotoxicity and metabolic diseases may lead to new treatment options.

#### Protein-Lipid Interactions

A more thorough understanding of perilipin 2’s function may come from in-depth research on how it interacts with different lipids and aids in lipid metabolism.

#### Comparative Genomics

The processes behind PLIN2’s conservation and divergence may be clarified by analyzing the genomic context and control of PLIN2 in other species.

#### Experimental 3D Structures

As PLIN2 experimental 3D structures become accessible, comparing them to predicted structures might reveal important information about how accurate predictions are.

